# Depression, GABA and age correlate with the plasma levels of inflammatory markers

**DOI:** 10.1101/689984

**Authors:** Amol K. Bhandage, Janet L. Cunningham, Zhe Jin, Qiujin Shen, Santiago Bongiovanni, Sergiy V. Korol, Mikaela Syk, Masood Kamali-Moghaddam, Lisa Ekselius, Bryndis Birnir

**Affiliations:** Department of Neuroscience, Uppsala University, BMC, Box 593, 751 24, Uppsala, Sweden; Department of Neuroscience, Psychiatry, Uppsala University, 751 85, Uppsala, Sweden; Department of Immunology, Genetics and Pathology, Science for Life laboratory, Uppsala University, 75108 Uppsala, Sweden

**Author notes:** contributed equally (Amol K. Bhandage, Janet L. Cunningham). **Corresponding author:** Bryndis Birnir, Department of Neuroscience, Uppsala University, 75124 Uppsala.

**Keywords:** depression, inflammation, inflammatory markers, psychiatric disorder, PBMC, age, CDCP1, CST5, GABA, GABA_A_ receptor, biomarker.

## Abstract

Immunomodulation is increasingly being recognised as a part of mental diseases. Here, we examined if levels of immunological protein markers altered with depression, age or by the inhibitory neurotransmitter gamma-aminobutyric acid (GABA). Analysis of plasma samples from patients with major depressive episode and control blood donors (CBD) revealed expression of 67 inflammatory markers. Thirteen of these markers had augmented levels in patients as compared to CBD. and 21 markers correlated with age of the patients, whereas 10 markers correlated with the age of CBD. Interestingly, CST5 and CDCP1 showed the strongest correlation with age in the patients and in the CBD, respectively. IL-18 was the only marker that correlated with the MADRS-S scores of the patients. Neuronal growth factors (NGFs) were significantly enhanced in plasma from the patients and so was the average plasma GABA concentration. GABA modulated release of seven cytokines in CD3^+^ stimulated peripheral blood mononuclear cells (PBMC) from the patients. The study reveals significant changes in plasma composition of small molecules during depression and identifies potential peripheral biomarkers of the disease.

## Introduction

Neurotransmitter signaling is well studied in the nervous system, where GABA is the main inhibitory transmitter [1]. Compelling evidence demonstrates that neurotransmitter signaling also takes place in the immune system [2-7]. The fact that a cross-talk occurs between the immune- and the nervous systems is not surprising. It may be required for normal brain functions and is probably essential for coordinated stress, emotional and behavioural responses [8]. Dysregulation of the immune system has furthermore been reported to be associated with psychiatric disorders, such as depression [8]. Pro-inflammatory cytokines can induce sickness behaviour that resembles major depressive disorder (MDD) and interferon-alpha (INF-α) treatment induces MDD in about 25% of cases, suggesting causal mechanisms [9, 10]. Pro-inflammatory markers such as IL-6, IL-1β, IFN-α, TNF-α and MCP-1/CCL2 are increased in the blood and cerebrospinal fluid (CSF) from patients with mood disorders compared to healthy controls, when assessed at baseline and also after exposure to stressors [11-13]. Inflammatory markers such as IL-6 and CRP are consistently found to be elevated in depression although the size of the effect is relatively small [14, 15]. Emerging evidence also indicates that antidepressants have immunomodulating effects and that inflammatory- and pro-inflammatory cytokines undermine the treatment response to conventional antidepressants [16-21]. Understanding the immunological changes in depression is important, as immunomodulation may be a possible therapy for some patients with depression.

In the brain, GABA is produced from glutamate in neuronal cells by the enzyme glutamic acid decarboxylase (GAD) [22]. Central nervous system (CNS) interstitial GABA and the human plasma GABA concentrations are expected to be in the submicromolar range [23-26], though the origin of GABA in blood is still being explored. Recently identified drainage system of the brain, the glymphatic system, indicates the brain as a significant source of the GABA present in blood [27]. The expression of GABA receptors subunits and activation of functional GABA_A_ receptors has been recorded in immune cells such as PBMCs, T cells, monocytes, dendritic cells and macrophages [7, 28]. Recently, we demonstrated that GABA inhibits secretion of a variety of inflammatory protein markers from PBMCs and T cells from healthy individuals and type 1 diabetes patients [7]. Nevertheless, the effects of GABA on secretion of cytokines/immunological markers from immune cells is still relatively unexplored.

Here, we analyzed the immunological markers in plasma, from CBD and patients with a major depressive episode, and how they altered with age. We further studied expression of the GABA signaling system components and effects of GABA treatment on the inflammatory markers profile of stimulated PBMCs from the patients. The results highlight augmented levels of immunological markers and the neurotransmitter GABA in the plasma of patients, together with altered GABA signaling in PBMCs from patients. The results are consistent with immunomodulatory effects of GABA during depression. Furthermore, the level of a number of inflammatory markers correlated with age for both groups.

## Results

Demographic data for the individuals (CBD:26; P:25) that participated in the study are shown in Table 1 and Supplementary Table 2. In total, 38 patients that met the criteria were selected for this study. Of these 38 patients, 7 patients chose not participate, while 6 individuals were found unable to provide informed consent due to cognitive symptoms. Thus, 25 patients were included in the study (Table 1). All the patients met the DSM IV criteria for current moderate to severe depressive episode and either major depressive disorder or bipolar disorder. They were all undergoing treatment of depression at the Department of General Psychiatry at Uppsala University Hospital, Sweden, at the time the samples were obtained. Eleven of the patients were prescribed benzodiazepines or “Z-drugs”, while three of the patients had both. None of the patients had a documented history of alcohol addiction or abuse disorder and none had consumed alcohol during the past week prior to the sampling. Two patients had received ECT during the past three months but none during the past month. Five patients have previously received ECT during their lifetime. Two patients had neurodevelopmental disorders but the physician evaluated them to be capable of judgment in giving consent. In three cases, the MINI interview could not be performed due to cognitive symptoms, one case developed psychotic symptoms with delusions and severe disorganized thinking 24 hours after giving consent, another patient presented severe concentrating difficulties and the last one did not consent to the interview due to fatigue. Diagnosis in these cases was made based on clinical records.

**Table 1.**
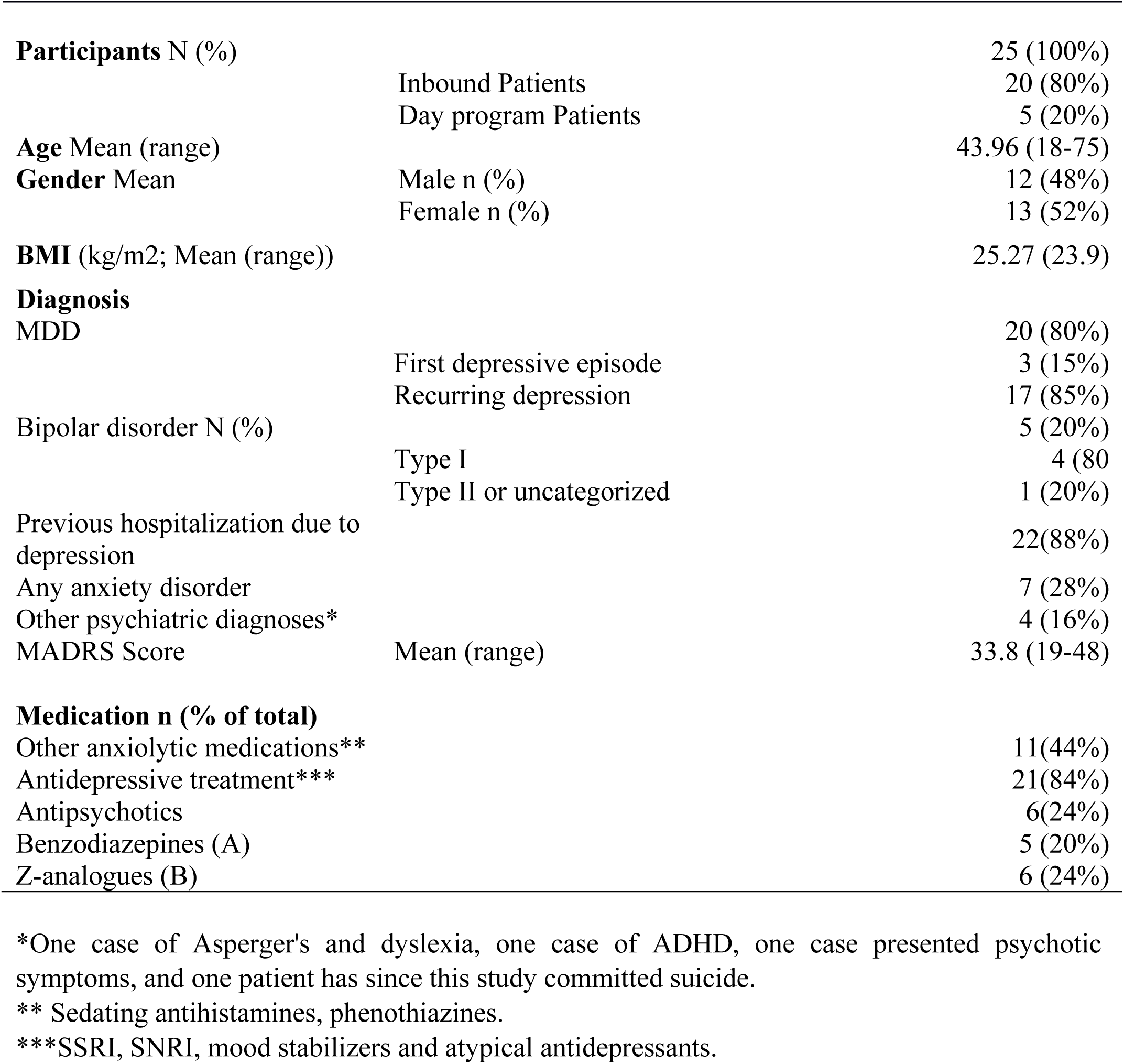
Patient’s characteristics.

### Inflammatory markers in plasma from patients and control donors

Immune cells release a large number of small proteins, collectively called inflammatory markers, which may have a protective function or act as pro-inflammatory molecules. We investigated whether the types of inflammatory markers in plasma differed between CBD and the patients. We measured the plasma levels of 92 inflammatory markers that are most commonly associated with inflammation using an Olink inflammation panel analyzed with multiplex PEA (Table S3) (http://www.olink.com/products/inflammation/#). The technology uses paired antibodies for the different inflammatory markers such as cytokines, growth factors, mitogens, chemotactic, soluble receptors and other pro-inflammatory molecules that allows comparison of the levels of the same marker in samples from e.g. CBD and the patients. However, the assay format does not support comparison of the absolute levels of one marker to another as the affinities of the antibodies for their cognate targets may vary. In plasma from both CBD and patient groups, 67 inflammatory markers out of 92 analyzed proteins were detectable with values above LOD (Fig. 1A; Table S4). Importantly, 13 inflammatory markers were significantly higher in plasma from the patients as compared to plasma from CBD (Fig. 1B).

**Figure 1.**
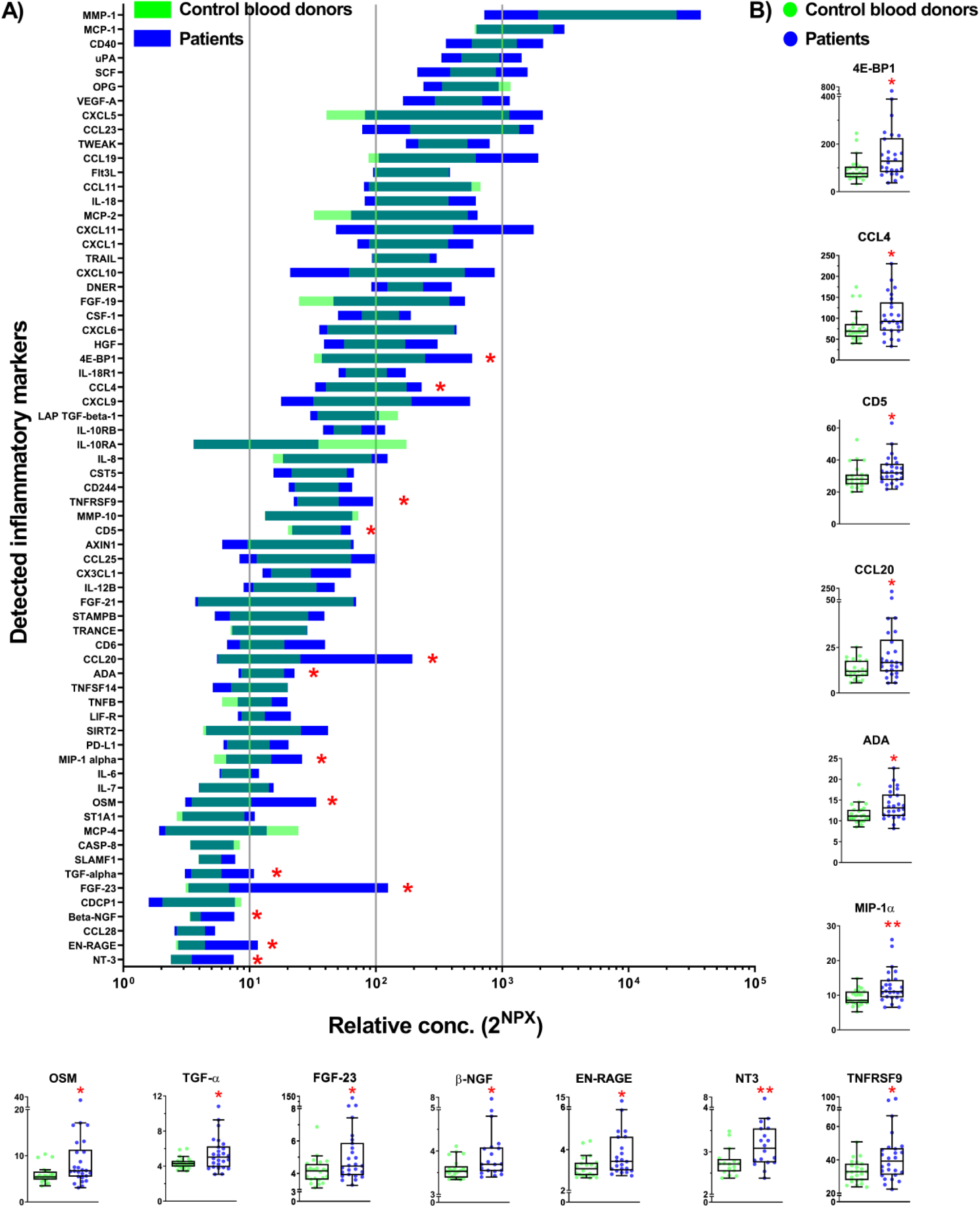
Inflammatory markers in plasma from control blood donors and patients. (A) Screening of 92 inflammatory markers (Table S3) in plasma samples from control blood donors (n=26) and patients (n=25) by Proseek Multiplex PEA inflammation panel I detects expression of 67 markers (Table S4). Data is presented by 2^NPX^ (Normalized Protein Expression) values as floating bars (minimum to maximum) arranged in descending order of mean expression level of inflammatory markers. (B) Inflammatory markers with significant change in the expression levels in the plasma of patients as compared to control blood donors. The differences between groups were assessed by nonparametric Kruskal–Wallis ANOVA on ranks with Dunn’s post hoc test. Data is shown as box and whiskers overlapped with scatter dot plot. * p < 0.05, ** p < 0.01.

### Effects of age on levels of inflammatory markers in plasma

Since the patients varied in age, we examined if there were any correlations between age and the level of the inflammatory markers in plasma from the two groups. Ten inflammatory markers correlated with age in CBD (Fig. 2A; Table S5) and 21 in the patients (Fig. 2B; Table S5). Six inflammatory markers, IL-8, CXCL9, HGF, VEGF-A, OPG, MMP-1, correlated with age in both groups and thus may reflect normal aging processes rather than disease. Another three inflammatory markers, TGF-α, EN-RAGE, OSM, only correlated with age in the patients but were also increased in plasma from patients as compared to CBD (Fig. 1B and 2B). The strongest correlation with age in CBD was observed for CDCP1, a molecule with a role in immune cells migration and chemotaxis [29-31], whereas in the patients, the strongest correlation was observed for CST5, a cysteine protease inhibitor which can also modulate gene transcription and protein expression [32, 33]. The inflammatory markers that varied with age can be grouped according to function and are shown in Figure 2C for the two groups. Inflammatory markers associated with activation of immune cell response and apoptosis were enhanced in plasma from the patients.

**Figure 2.**
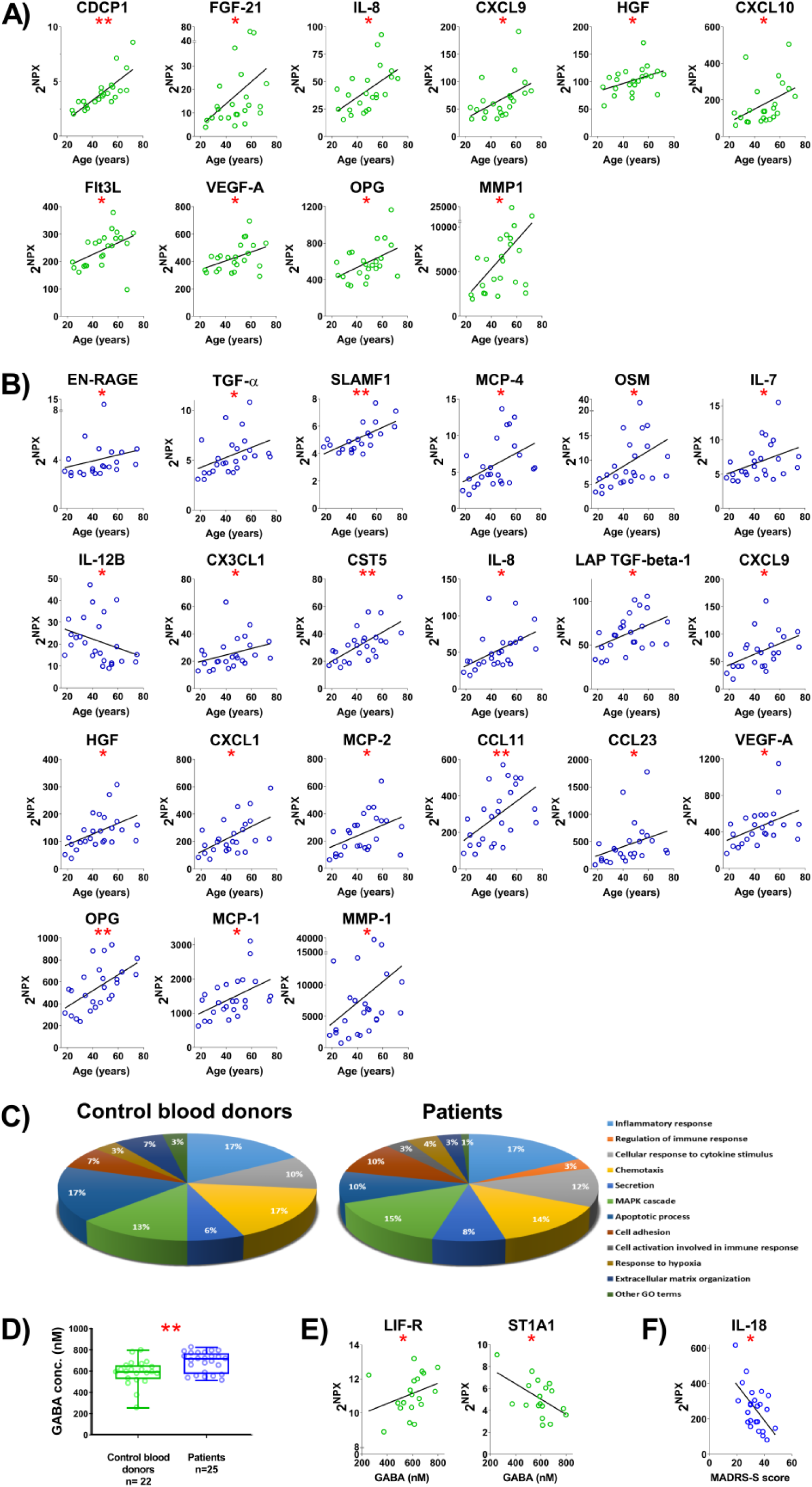
Inflammatory markers correlation with age and GABA in control blood donors and patients. Correlation between inflammatory marker levels in plasma samples and age of (A) control blood donors and (B) patients. Only inflammatory markers with significant correlation are shown. (C) Classification based on the cellular functions of markers that were significantly correlated with age of control blood donors (10 inflammatory markers) and patients (21 inflammatory markers). (D) Quantification of GABA levels in plasma samples from control blood donors and patients. (E) Correlation between inflammatory marker levels and GABA levels in plasma samples control blood donors. (F) Correlation between inflammatory marker levels and MADRS score in plasma samples control blood donors. The correlation between inflammatory markers and demographic factors was accessed using non-parametric Spearman rank correlation. To reduce the risk of false discoveries caused by multiple testing, the Benjamini-Hochberg false discovery rate method was used. Rho values and p values of correlation statistics are provided in Table S5. * p < 0.05, ** p < 0.01, *** p < 0.001.

### The GABA concentration in plasma and correlation of GABA or MADRS-S score with inflammatory markers

Since GABA is exclusively generated within the body, and is the main inhibitory neurotransmitter in the brain, we examined if the GABA concentration in plasma varied between CBD and the patients (Fig. 2D). The results showed that the GABA concentration ranged from 253 to 824 nM in the two groups and revealed somewhat increased plasma GABA concentration in the patients, resulting in a significantly higher average plasma concentration (CBD: 586±25 nM; PD: 683±19 nM; p=0.003). In general, the GABA levels in plasma did not correlate with the inflammatory marker levels in plasma with the exception of LIF-R, ST1A1 in the CBD (Fig. 2E, Table S5), neither with the age of the CBD or the patients nor with the MADRS-S score or the BMI of the patients. A posthoc analysis with t-test showed elevated levels of GABA in the patients with benzodiazepines and/or Z-drugs when compared to the remaining patients (t-value −2.354, p-value 0.037). Importantly, most of these patients were also treated with other medication with potential for influencing GABA. None of the markers correlated with BMI, whereas IL-18 was the only marker that correlated with the MADRS-S score of the patients (Fig. 2F; Table S5, R-value −0.4832, p-value 0.017).

### The GABA signaling system is altered in PBMCs from the patients

GABA can activate two types of receptors in the plasma membrane of cells, the GABA_A_ receptors that are chloride ion channels opened by GABA and the G-protein-coupled GABA_B_ receptor [1, 34, 35]. The GABA_A_ receptors are homo- or hetero-pentamers formed by a selection of subunits from 19 known isoforms (α1-6, β1-3, γ1-3, δ, ε θ, π, ρ1-3) [35]. In contrast, the GABA_B_ receptor is normally formed as a dimer of the two isoforms identified to-date [34, 36]. The ρ2 subunit was the only GABA_A_ subunit, which was expressed in PBMCs from most of the CBD and the patients (Fig. 3A, Table 2). The expression level was similar in both groups and could indicate the formation of homomeric ρ2 GABA_A_ receptors in the cells. Approximately 30-40% of the CBD also express the β1, δ and ε subunits, while the expression of these subunits was less frequent in the patients (Table 2). Other GABA_A_ subunits were expressed only infrequently in both groups (Table 2). Only one GABA_B_ subunit was expressed in both patients and CBD indicating that the traditional GABA_B_ receptors may not be formed in the PBMCs (Table 2).

**Figure 3.**
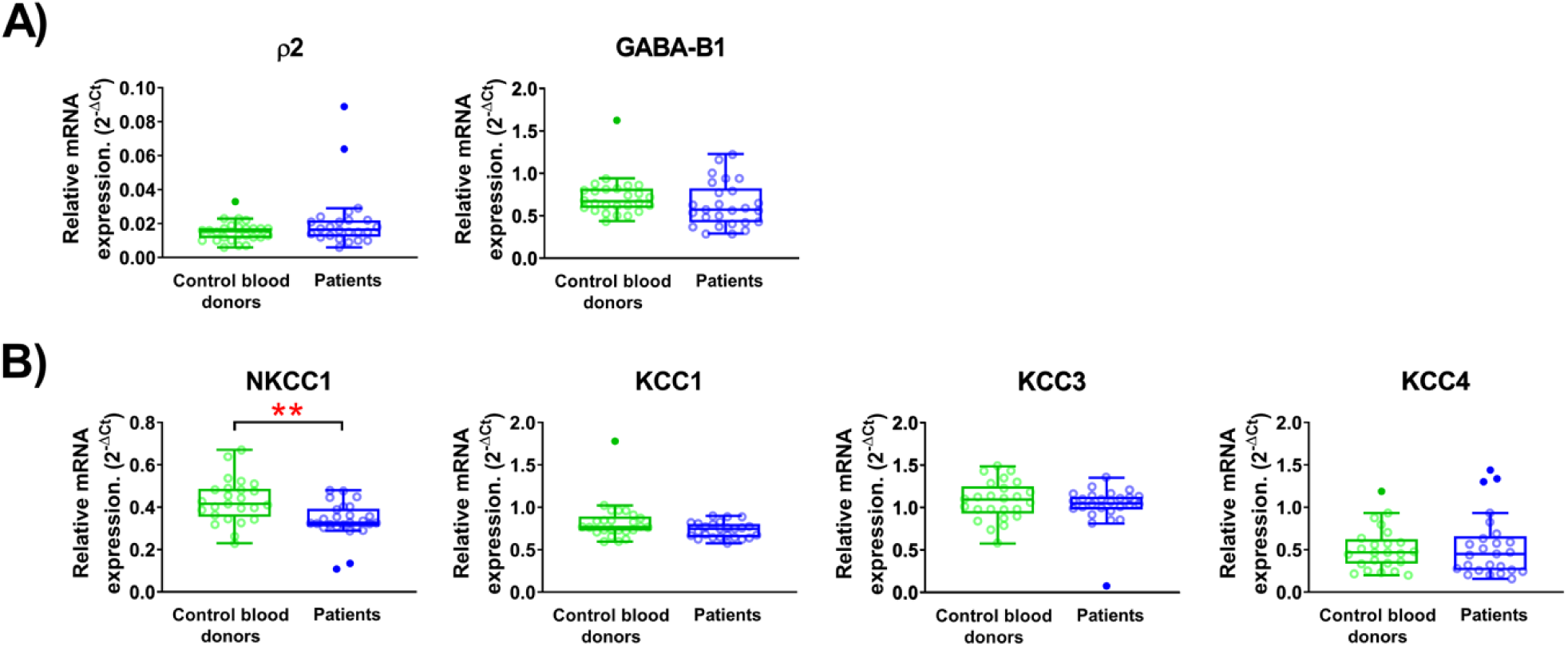
The relative mRNA expression in PBMCs from control blood donors and patients. (A) GABA_A_ receptor subunit ρ2 and GABA_B_ receptor subunit 1 (B) chloride co-transporters, NKCC1, KCC1, KCC3, KCC4. Data are shown as box and whiskers overlapped with scatter dot plot. The outliers were detected using Tukey’s test (with 1.5 times +/− IQR, inter quartile range) and are shown with filled circles. Normality of data was assessed by Shapiro-Wilk normality test (Table S8). ** p < 0.01.

**Table 2.** The percentage of samples expressing the particular mRNA.

|  | CBD | Patients |
| --- | --- | --- |
| **GABA<sub>A</sub> receptor subunits** |  |  |
| GABRA1 (α1) | 0 | 0 |
| GABRA2 (α2) | 0 | 0 |
| GABRA3 (α3) | 3.8 | 4 |
| GABRA4 (α4) | 15.4 | 8 |
| GABRA5 (α5) | 19.2 | 4 |
| GABRA6 (α6) | 15.4 | 16 |
| GABRB1 (β1) | 30.8 | 8 |
| GABRB2 (β2) | 38.5 | 36 |
| GABRB3 (β3) | 0 | 0 |
| GABRG1 (γ1) | 0 | 4 |
| GABRG2 (γ2) | 0 | 0 |
| GABRG3 (γ3) | 0 | 0 |
| GABRD ($\delta$) | 34.6 | 12 |
| GABRE ($\epsilon$) | 42.3 | 20 |
| GABRQ ($\theta$) | 0 | 0 |
| GABRP ($\pi$) | 3.8 | 4 |
| GABRR1 (ρ1) | 0 | 0 |
| GABRR2 (ρ2) | 100 | 96 |
| GABRR3 (ρ3) | 0 | 12 |
| **GABA<sub>B</sub> receptor subunits** |  |  |
| GABBR1 (GABA-B1) | 100 | 100 |
| GABBR2 (GABA-B2) | 0 | 0 |
| **Chloride transporter** |  |  |
| SLC12A2 (NKCC1) | 100 | 100 |
| SLC12A1 (NKCC2) | 0 | 0 |
| SLC12A4 (KCC1) | 100 | 100 |
| SLC12A5 (KCC2) | 0 | 0 |
| SLC12A6 (KCC3) | 100 | 100 |
| SLC12A7 (KCC4) | 96 | 100 |

Total of 51 PBMC samples were examined, 26 from control blood donors and 25 from patient

The strength of GABA_A_ receptor signaling depends, in part, on the chloride gradient across the cell membrane. We, therefore, examined if the expression of chloride transporters that regulate the intracellular chloride concentration differed between the CBD and the patients (Fig. 3B). The NKCC1 transporter that transports chloride ions into the cells was significantly down-regulated in the PBMCs from the patients. The other three transporters, that move Cl^−^ out of the cells, were also expressed in the majority of the samples but at similar levels in both groups.

### GABA regulates release of inflammatory markers from stimulated PBMCs from the patients

Since the levels of inflammatory markers in plasma differed between the two groups and GABA has been reported to be immunomodulatory [7], we examined if GABA affected release of inflammatory markers from the patients’ PBMCs. The culture media of anti-CD3 stimulated PBMCs were assessed using the same Olink Inflammation protein panel composed of 92 inflammatory markers used for analysis of the plasma samples described above. We also examined if GABA at concentrations of 100 and 500 nM regulated secretion of specific inflammatory markers.

A total of 59 different inflammatory markers were detected in the culture media from both non-stimulated and the anti-CD3 stimulated PBMCs (Fig. 4A; Table S6), and additional nine markers (IL-2, IL-4, IL-5, IL-13, IL-2RB, IL-10RA, IL-15RA, PD-L1 and SLAM-F1) in the media from the stimulated cells. The majority of the inflammatory markers were secreted at a higher level in the stimulated PBMCs (Fig. 4A). Only VEGF-A was secreted at a significantly higher level in the resting state of the cells (Fig. 4A). From the stimulated PBMCs, 51 inflammatory markers were the same as those identified in the plasma samples, while the remaining 17, including INF-γ, TNF-α and IL-1α, were not detected in the plasma samples. We further examined if treatment with GABA altered the release of inflammatory markers from the stimulated PBMCs as compared to non-treated PBMCs. When 100 nM GABA was applied, there was a significant increase in secretion of 2 inflammatory markers, AXlN1 and TNFRSF9, and a decrease of another two, VEGF-A and IL-1α (Fig. 4B, Table S7). With 500 nM GABA treatment, secretion of 4 inflammatory markers was significantly decreased from the stimulated PBMCs; CD244, IL-13, HGF and VEGF-A (Fig. 4C; Table S7). Of the inflammatory markers regulated by GABA, HGF and VEGF-A, increased with age in both, CBD and the patients (Fig. 1B). Interestingly, of these seven inflammatory markers, only VEGF-A was modulated by both concentrations of GABA.

**Figure 4.**
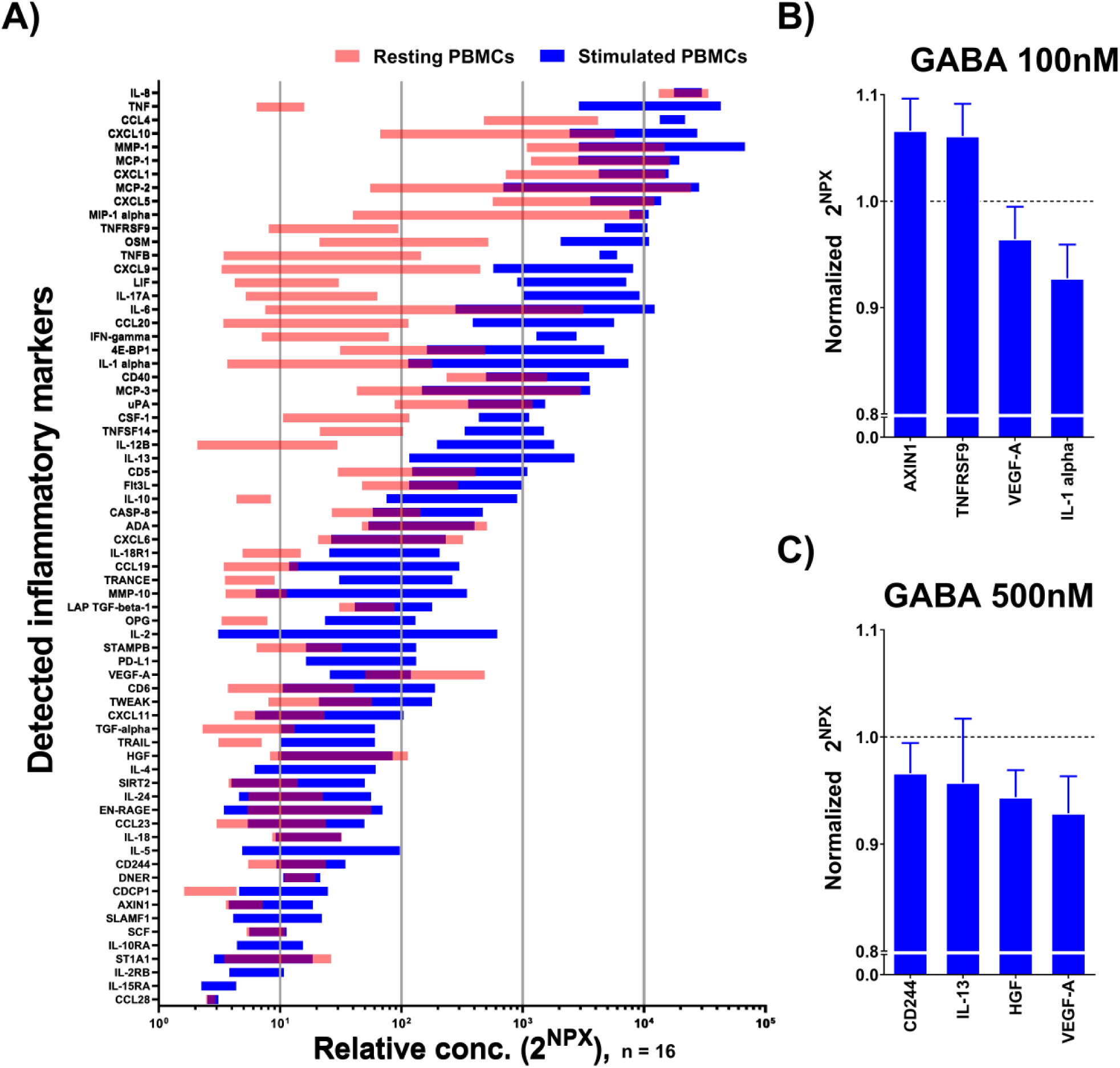
Identification of cytokines released from patients PBMCs and effects of GABA treatment on the inflammatory markers secretion. (A) Screening of 92 inflammatory markers (Table S3) in PBMCs media from patients by Proseek Multiplex PEA inflammation panel I revealed expression of 59 (light red) and 68 (blue) inflammatory markers from non-stimulated and anti-CD3 stimulated PBMCs, respectively (Table S6). Data is represented by 2^NPX^ values as floating bars (minimum to maximum) arranged in descending order of mean expression level of the inflammatory markers. (B-C) Expression of inflammatory markers that are significantly affected by (B) GABA 100 nM and (C) GABA 500 nM treatment of anti-CD3 stimulated PBMCs from patients. Data is represented by 2^NPX^ values normalized to controls as a bar graph with mean ± SEM. Mean values with SEM and p values are provided in Table S7. The differences between groups were assessed by nonparametric Kruskal–Wallis ANOVA on ranks with Dunn’s post hoc test (Table S7). p < 0.05 for (B) and (C).

## Discussion

This study examined peripheral inflammatory markers and the immunoregulatory effects of GABA and GABA signaling in PBMCs from patients with a major depressive episode (Fig. 5). The results identified thirteen inflammatory markers that were upregulated in plasma from the patients. Consistent with other studies, a number of inflammatory markers correlated with age for both CBD and the patients, but the most prominent age-associated marker differed for the two groups, being CDCP1 for the CBD and CST5 for the patients. The main GABA_A_ receptor in the PBMCs was unchanged in the patients PBMCs but the NKCC1 transporter was down-regulated. Physiological concentrations of GABA modulated secretion of inflammatory markers from the patient’s immune cells. The results support an immunoregulatory role of GABA-activated GABA_A_ receptor signaling in the PBMCs.

**Figure 5.**
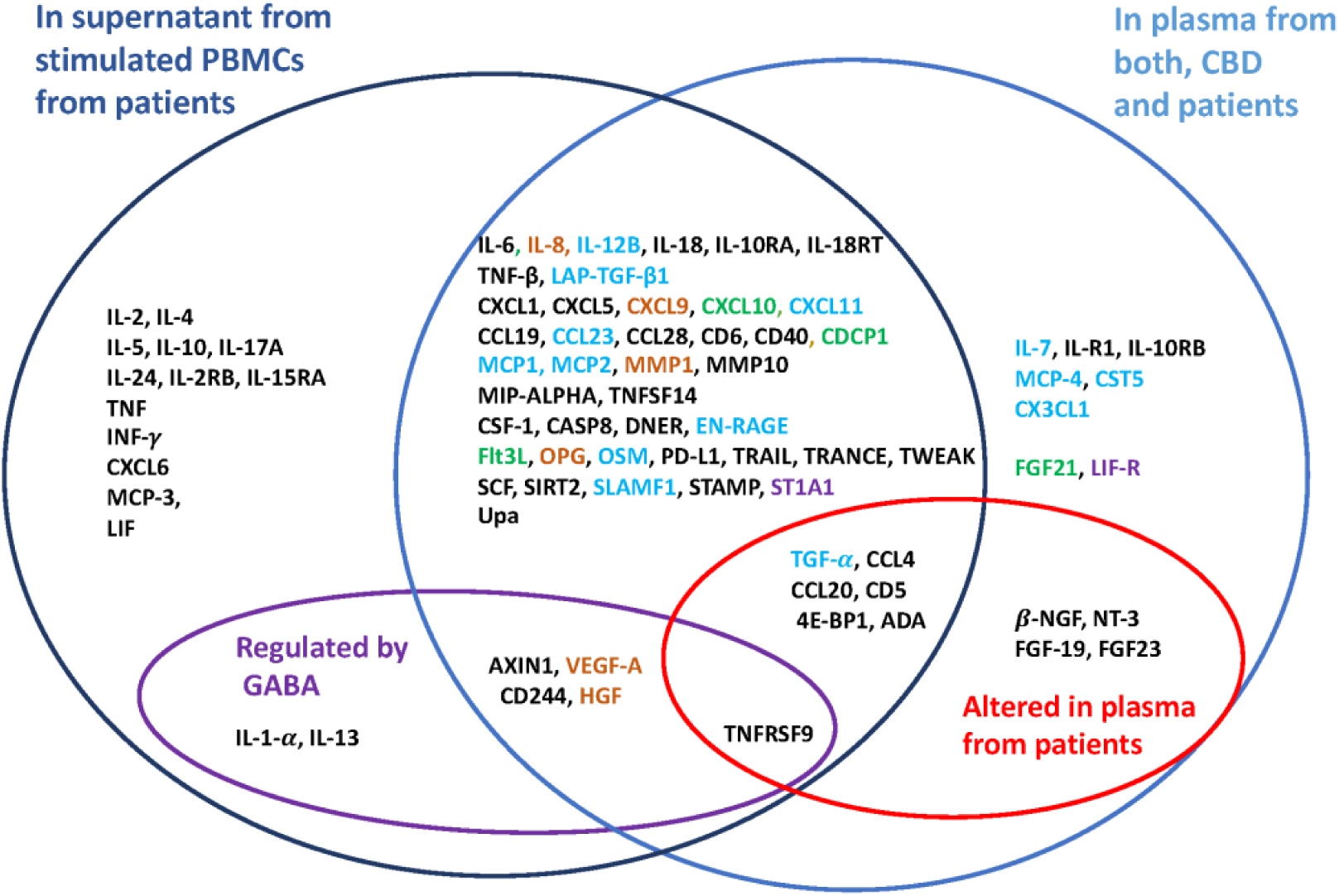
Inflammatory markers in plasma or released from stimulated PBMCs *in vitro*. *Dark blue circle:* Inflammatory markers detected in supernatants from anti-CD3 antibody stimulated PBMCs from patients. *Light blue circle:* Inflammatory markers detected in plasma samples from control blood donors (CBD) and patients (P). *Violet circle:* Inflammatory markers regulated by GABA in PBMCs from (P). *Red circle:* Inflammatory markers altered in plasma of (P) compared to (CBD). *Blue:* Inflammatory markers that correlated with age in only (P), *green:* Inflammatory markers that correlated with age in only (CBD), *brown:* Inflammatory markers that correlated with age in both (P) and (CBD), *violet:* Inflammatory markers that correlated with GABA concentration in blood from (CBD).

The inflammatory environment is thought to be altered in many psychiatric disorders [8, 9, 12]. The results in this report showed clear differences at the systemic level with the altered plasma concentrations of specific inflammatory markers in the patients (Fig. 5). Inflammatory markers released from stimulated PBMCs from the patients identified additional markers that may be significant as auto- or paracrine signals. In the present study, the patients had increased levels of NGFs and neurotropin-3 (NT3) as compared to CBD and, notably, NGFs were not released from the simulated PBMCs. Low levels of NGFs have been implicated in the pathogenesis of depression and observed in patients with depressive disorders [37, 38]. β-NGF and NT3 are the NGFs that increase viability, growth and development of neurons and may supress inflammation [39]. It is possible that the increase in NGFs levels in the patients is related to improvements in response to the antidepressants. This is in accordance with studies in animal models where increased expression/concentration levels of NT3 and NGF was observed in a number of brain regions and/or in serum in response to treatments with antidepressants [40-43]. Interestingly, a recent study of inflammatory markers altered in plasma from patients with the autoimmune disease type 1 diabetic (T1D) or secreted by stimulated T1D PBMCs, identified many of the same inflammatory markers as in this study [7]. How small molecules from the brain reach the circulation is still being explored but they may diffuse by volume transmission from the brain to the blood [27] by the route of the active glymphatic system.

A number of studies have reported alteration in inflammatory markers with medical treatment. Recent meta-analysis studies have shown decreased plasma levels of IL-6, IL-10, IL-1β, TNF-α, p11, IFN-γ and CCL2 after treatment with antidepressants, including SSRI and SNRI [18, 44-46]. Reduced mRNA expression of IL-6, IL-1b and MIF in leukocytes after treatment of patients with antidepressants, escitalopram or nortriptyline, has been reported [21]. Additionally, high levels of VEGF-A mRNA in whole blood from the patients with depression were reported although the VEGF protein levels in the plasma were not affected [47]. Further, increased IFNγ/IL-10 ratio and changes in CCL11 and IFNγ with antidepressant treatments has been reported [48, 49] but lithium augmentation of antidepressants had no effect on the inflammatory markers in MDD [50].

Several inflammatory markers, that are often reported to be altered in patients with MDD, did not differ significantly between the patients in this study and CBD, e.g. IL-6. A part of the explanation may potentially be that the antidepressant treatment normalized the levels of some of the inflammatory markers. However, MIP-1α (i.e. CCL3), CCL4 and CCL20, the macrophage-released pro-inflammatory molecules, were significantly elevated in the patients consistent with previous MDD reports [51, 52]. IL-18 was the only inflammatory marker that correlated with the MADRS-S score and is consistent with other studies of inflammatory markers and depression [53-56]. Many inflammatory markers correlated with age in both CBD and the patients in concordance with previous studies [57-60]. Our study corroborates that CDCP shows the strongest correlation with age in healthy individuals [59] whereas CST5 (cystatin D) with age in the patients. CST5 is an inhibitor of lysosomal and secreted cysteine proteases, but can also locate to the nucleus and modify gene transcription [33]. Interestingly, CST5 is an ultra-early biomarker of traumatic brain injury [32].

The average plasma levels of GABA were increased in the patients as compared to age and sex matched CBD GABA levels, but there was, nevertheless, a narrow range of values and considerable overlap of sample GABA concentrations observed between the two groups. Circulating levels of GABA are much lower than those found in the synaptic cleft and instead comparable to levels activating high affinity extrasynaptic receptors, generating small amplitude but long lasting currents [1]. In previous studies where plasma GABA levels have been measured in samples from patients with psychiatric disorders and a control group, the results have varied [26, 61-66]. Since the majority of the patients in this study were being treated (medicines e.g. benzodiazepines, Li^+^, valproate or antipsychotic medication), it is possible that the difference observed in the GABA plasma concentration is related to the effect of the medications [65, 67].

We and others have shown that immune cells can be regulated by GABA. GABA can e.g. decrease proliferation of T cells, reduce inflammation in experimental autoimmune encephalomyelitis, decrease cytokine secretion from T cells and modulate mobility of infected dendritic cells [3-5, 7, 68, 69]. The PBMCs express genes for the diverse components of the GABA signaling system including the GABA_A_ receptors and the chloride transporters [7] and respond to GABA by activating GABA_A_ receptors channels [4, 7]. NKCC1, the transporter that maintains high intracellular chloride in the cells, was down-regulated in PMBCs from the patients implying decreased strength of GABA signaling in the cells. The alteration in the chloride gradient across the plasma membrane is potentially partially compensated for by the somewhat increased GABA concentration in the plasma. Similar down-regulation of NKCC1 has been observed in cells from healthy and depressed pregnant women [28] and T1D patients [7]. This observation thus indicates a general shift of immune regulation by altered GABA signaling, rather than a decrease specifically associated with depression. Another possibility is that reduced expression of NKCC1 is a trait conferring vulnerability for depression.

This study has several limitations. First, the patients were recruited in a naturalistic setting and have different combinations of medications that may influence GABA signaling. The careful selection of patients has, however, reduced the effects of other confounders, such as alcohol use and recent electroconvulsive therapy. Secondly, patients with depression are in different stages of the disease process. Thirdly, as this is a pilot study, we have not done correction for multiple testing and the findings must be validated in new cohorts. Finally, here the control blood was obtained from the hospital blood-central facility and donors were only matched for sex and age and were not evaluated in terms of mental health. A strength of this study is the inclusion of patients with severe depressive states which increases the likelihood that these results are relevant for a clinical psychiatric population.

The study shows that significant changes take place in the immune system during depression andidentifies molecules and mechanism important for immunomodulation in depression.

(Fig. 5). Levels of several inflammatory molecules including NGFs, IL-18 and CST5 were altered in plasma of patients with depression. In PBMCs from the patients GABA modulated release of cytokines. The average GABA level in the plasma from patients was increased whereas the NKCC1 expression in the PBMCs was decreased. Together the results suggest altered GABA signaling during depression. Future studies are required to understand if specific subpopulations of immune cells are involved in mental illness and then, how they are regulated. Further, the link between concentrations of small molecules like GABA, NGFs and CST5 in plasma and brain functions needs to be explored.

## 2. Materials and Methods

### Study individuals, ethical permits and blood samples

Psychiatric illness was diagnosed using the International Neuropsychiatric Interview (M.I.N.I. 6.0) and the Diagnostic and Statistical Manual of Mental Disorders (DSM)-IV criteria. The interviews were conducted by two resident physicians in psychiatry and a specialized nurse in psychiatry at the clinic [70]. Patients were recruited at the Department of General Psychiatry at Uppsala University Hospital, Sweden, and the inclusion criteria for this study was that they met the DSM IV criteria for current moderate to severe depressive episode and either major depressive disorder or bipolar disorder at the time of blood sample collection. Depression severity was assessed using the self-rating version of Montgomery Åsberg depression rating scale (MADRS) [71, 72]. It is a 10-item clinician-rated scale measuring severity of depressive symptoms including the following items: reported sadness, inner tension, reduced sleep, reduced appetite, concentration difficulties, lassitude, inability to feel, pessimism and suicidal thoughts. The items are rated on a Likert scale from 0 to 6 and the total score ranges from 0 to 60. Higher scores indicate a greater severity. The study was approved by the Regional Ethics Committee in Uppsala (D.nr 2014/148 2014-06-12 and 2015-11-02), and all participants provided written consent. Twenty-five psychiatric patients with a major depressive episode participated in the study. Venous blood samples from the patients (P) were collected in EDTA tubes and used to isolate plasma and PBMCs. Twenty-six control blood samples from blood donors (CBD) were obtained at the blood center at Uppsala University Hospital.

### Plasma and PBMC isolation from blood samples

Plasma and PBMCs were isolated from freshly drawn derived blood samples as previously described [28]. The plasma was isolated by centrifugation at 3,600 rpm for 10 min at 4 °C and immediately frozen at −80 °C. PBMCs were prepared by first diluting the blood samples in equal volume of MACS buffer (Miltenyi Biotec, Madrid, Spain), and layered on Ficoll-paque plus (Sigma-Aldrich, Hamburg, Germany). Briefly, the samples were then subjected to density gradient centrifugation at 400 g for 30 min at room temperature. The PBMCs were carefully withdrawn and washed twice in MACS buffer. A portion of purified PBMCs was saved in RNAlater (Sigma-Aldrich) at −80 °C for mRNA extraction for qPCR experiments, and the remaining portion was used for analysis of cell culture supernatants by multiplex proximity extension assay (PEA).

### Multiplex PEA for inflammatory marker measurements

Plasma samples, and culture media supernatants were analyzed using multiplex PEA with a panel of 92 inflammation-related proteins (multiplex Inflammation I^96×96^, Olink Proteomics^TM^, Uppsala, Sweden) as previously described [73]. In brief, 1 µl of sample or negative control was incubated overnight at 4 °C with a panel of oligonucleotides-conjugated antibodies, where each target protein can be recognized by a pair of antibodies. This binding brings the attached oligonucleotides in close proximity, allowing them to hybridize to each other and subsequently extended via enzymatic DNA polymerization, creating DNA amplicons, which are quantified using microfluidic-based quantitative real-time PCR system (Fluidigm, San Francisco, CA, USA). To even out intra-plate variations, the raw quantification cycle (Cq) values were normalized against spiked-in controls and negative controls to achieve normalized protein expression (NPX) values. NPX is an arbitrary value in log_2_ scale, where an increase of a unit corresponds to a two-fold increase of protein concentration. These NPX data were then converted to linear data, using the formula 2^NPX^, prior to further statistical analysis. Each protein has its own limit of detection (LOD) defined as the NPX of background plus three times standard deviations. The multiplex PEA is reported to have a sensitivity in subpicomolar and a broad dynamic range. The technical performances including LOD, dynamic range, etc., for all the proteins included in the panel is available at the manufacturer’s homepage: https://www.olink.com/data-you-can-trust/publications. Proteins with levels below LOD were excluded from further data analysis.

### Determination of GABA concentration

Plasma samples were thawed and the levels of GABA were measured using an ELISA kit (LDN Labor Diagnostika Nord, Nordhorn, Germany) as per manufacturer’s guidelines [69]. In brief, plasma samples and standards provided in the kit were extracted on extraction plate, derivatized using equalizing reagent and subjected standard competitive ELISA in GABA coated microtiter strips. The absorbance of the solution in the wells was read at 450 nm within 10 min using Multiskan MS plate reader (Labsystems, Vantaa, Finland). Optical density was used to calculate the GABA concentration using a standard curve.

### Real-Time Quantitative Reverse Transcription PCR

The PBMC samples collected from 26 CBD and 25 patients were subjected to total mRNA extraction using RNA/DNA/Protein Purification Plus Kit (Norgen Biotek, Ontario, Canada). RNA concentration was measured using Nanodrop (Nanodrop Technologies, Thermo Scientific, Inc., Wilmington, DE, USA). Further, 1.5 μg RNA was treated with 0.6 U DNAase I (Roche, Basel, Switzerland) for 30 min at 37 °C, with 8 mM EDTA for 10 min at 75 °C and then converted to cDNA using Superscript IV reverse transcriptase (Invitrogen, Stockholm, Sweden) in a 20 μl reaction. Reverse transcriptase negative reaction was also carried out in order to confirm the absence of genomic DNA contamination. The gene-specific primer pairs are listed in Table S1. The PCR amplification was performed using the ABI PRISM 7900 HT Sequence Detection System (Applied Biosystems) with an initial denaturation step of 5 min at 95 °C, followed by 45 cycles of 95 °C for 15 s, 60 °C for 30 s and 72 °C for 1 min.

### PBMC supernatants for multiplex PEA

The cells were suspended in complete medium (RPMI 1640 supplemented with 2 mM glutamine, 25 mM HEPES, 10% heat inactivated fetal bovine serum, 100 U/ml penicillin, 10 mg/ml streptomycin, 5 μM β-mercaptoethanol) in a concentration of 1 million cells per ml. The cells (100 µl = 100000 cells) were added in triplicate for each experimental group to the 96-well plates pre-coated with 3 μg/ml anti-CD3 antibody (clone HIT3a, BD Biosciences) for 3-5 h at 37 °C. The cells were then incubated in presence or absence of GABA at the relevant concentration for 72 h at 37 °C (95% O_2_, 5% CO_2_) and supernatant culture media were collected, centrifuged to remove cellular debris and stored at −80 °C for the analysis of inflammatory markers using the multiplex PEA as mentioned above.

### Statistical Analysis

Statistical analysis and data mining were performed using Statistica 12 (StatSoft Scandinavia, Uppsala, Sweden) and GraphPad Prism 7 (La Jolla, CA, USA). The statistical tests were performed after omitting outliers identified by the Tukey test. The differences between groups were assessed by nonparametric Kruskal–Wallis ANOVA on ranks with Dunn’s post hoc test. The contingency of sex equality between the two groups was accessed by Fisher’s exact test and age was accessed by non-parametric Mann-Whitney test. The correlation between inflammatory markers and demographic factors was accessed using non-parametric Spearman rank correlation. To reduce the risk of false discoveries caused by multiple testing, the Benjamini-Hochberg false discovery rate method was used [74]. The significance level was set to p < 0.05.

## Acknowledgements

We thank the patients who participated in this study and the people who have donated blood at the Uppsala Academic Hospital blood central. We thank Karin Nygren for help with isolation of PBMCs and Johan Lundgren and Svante Berg for help in collecting blood samples from patients. The work in this study was supported by the Swedish Research Council, the Märta och Nicke Nasvells fund; The Swedish Society of Medicine; GastricGlycoExplorer funded by Marie Curie ITN and Medical Training and Research Agreement (ALF) Funds from Uppsala University Hospital.

## Author Contributions

B.B. and L.E. conceived the study, A.K.B. executed most experiments, Q.S. participated in PEA experiments A.K.B., Z.J., Q.S., S.K., J.C., M.K.M. and B.B. analysed data. J.C, and S.B. recruited, diagnosed and collected clinical data and blood samples from the patients, B.B. wrote the first draft of the manuscript that was then critically reviewed by A.K.B, J.C., L.E., M.S., Z.J., M.K.M. and then the other co-workers.

## SUPPLEMENTARY INFORMATION

**Table S1:**
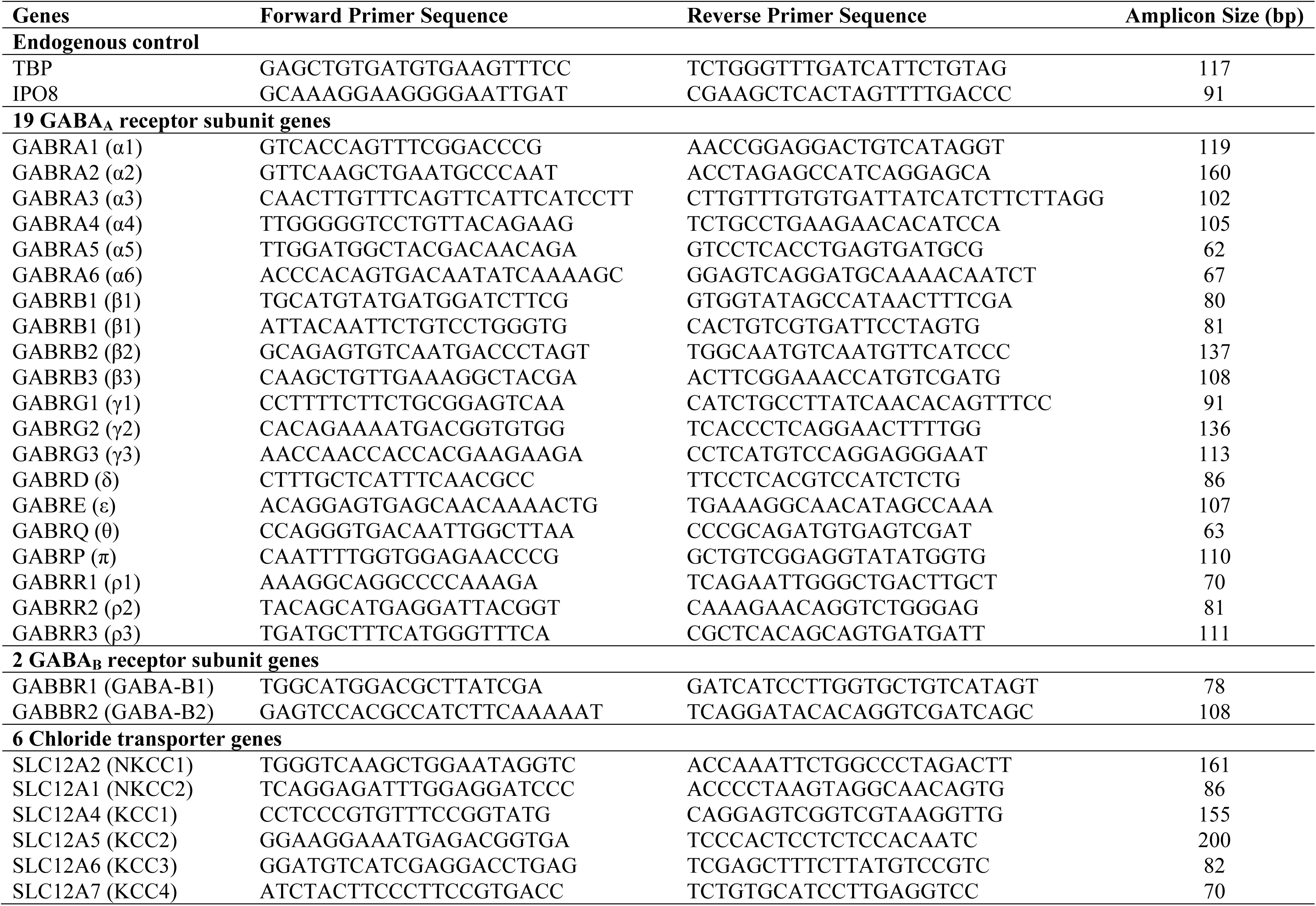
Primers for RT-qPCR.

**Table S2:** Demographic characteristic of all study participants.

| Control blood donors |  | Patient |  |  |
| --- | --- | --- | --- | --- |
| Age (Years) | Sex | Age (Years) | Sex | MADRS-S score |
| 55 | Woman | 75 | Man | 27 |
| 52 | Man | 55 | Woman | 48 |
| 34 | Woman | 42 | Man | 19 |
| 47 | Man | 38 | Woman | 46 |
| 47 | Man | 63 | Woman | 35 |
| 55 | Woman | 23 | Man | 21 |
| 48 | Man | 46 | Woman | 34 |
| 67 | Woman | 23 | Woman | 39 |
| 56 | Man | 40 | Woman | 30 |
| 24 | Man | 30 | Man | 36 |
| 58 | Woman | 21 | Woman | 28 |
| 67 | Woman | 45 | Woman | 33 |
| 59 | Man | 59 | Man | 42 |
| 36 | Woman | 53 | Woman | 30 |
| 33 | Man | 55 | Man | 43 |
| 35 | Woman | 49 | Woman | 30 |
| 42 | Woman | 33 | Man | 38 |
| 72 | Man | 74 | Man | 42 |
| 45 | Man | 48 | Man | 35 |
| 62 | Woman | 34 | Woman | 30 |
| 49 | Woman | 27 | Man | 39 |
| 29 | Woman | 49 | Man | 31 |
| 25 | Woman | 18 | Man | 28 |
| 44 | Woman | 40 | Woman | 24 |
| 24 | Man | 59 | Woman | 37 |
| 25 | Man |  |  |  |

The contingency of sex equality between the two groups was accessed by Fisher's exact test ( $p = 0.99$ ) and age was accessed by non-parametric Mann-Whitney test ( $p = 0.61$ ).

**Table S3:**
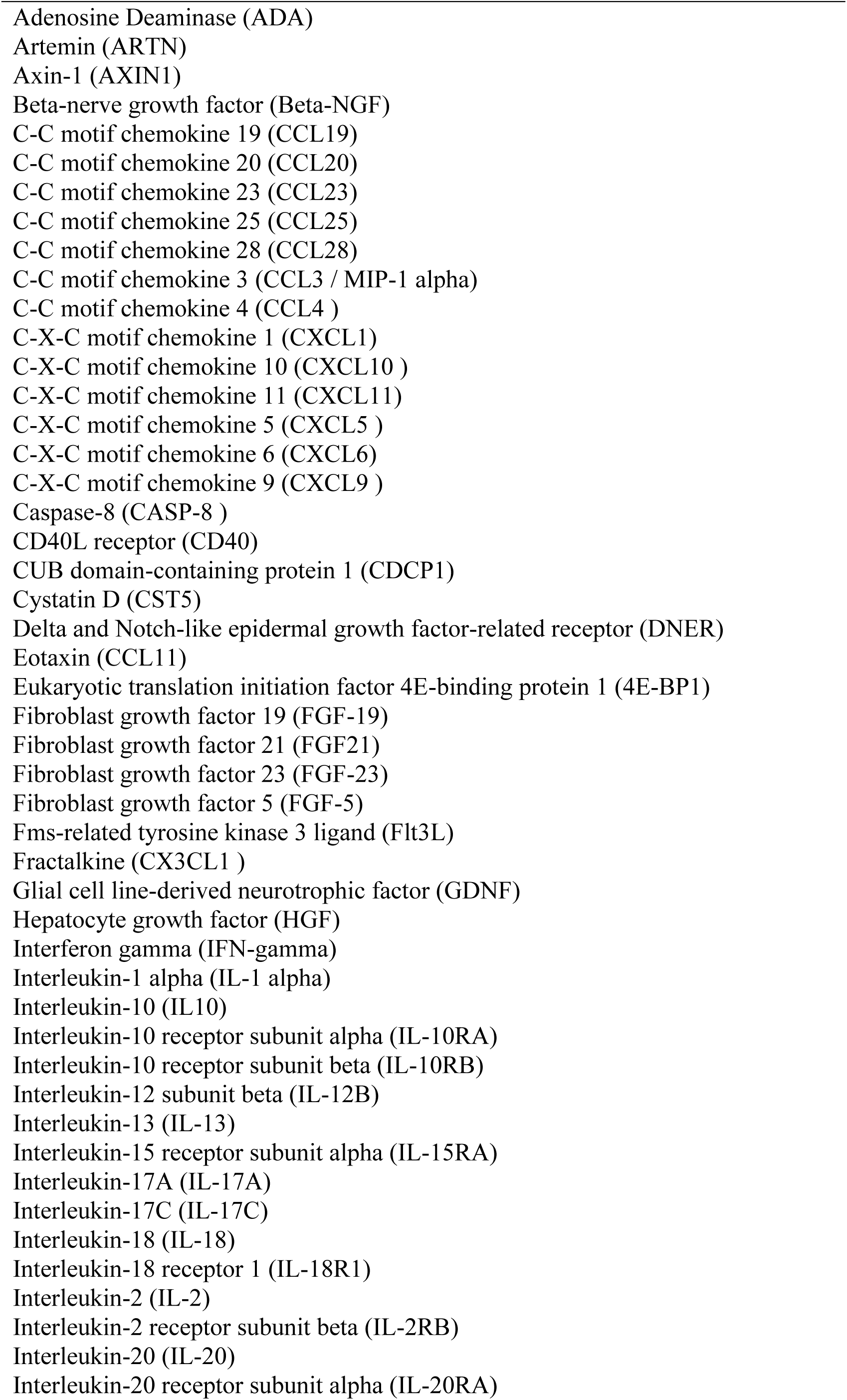
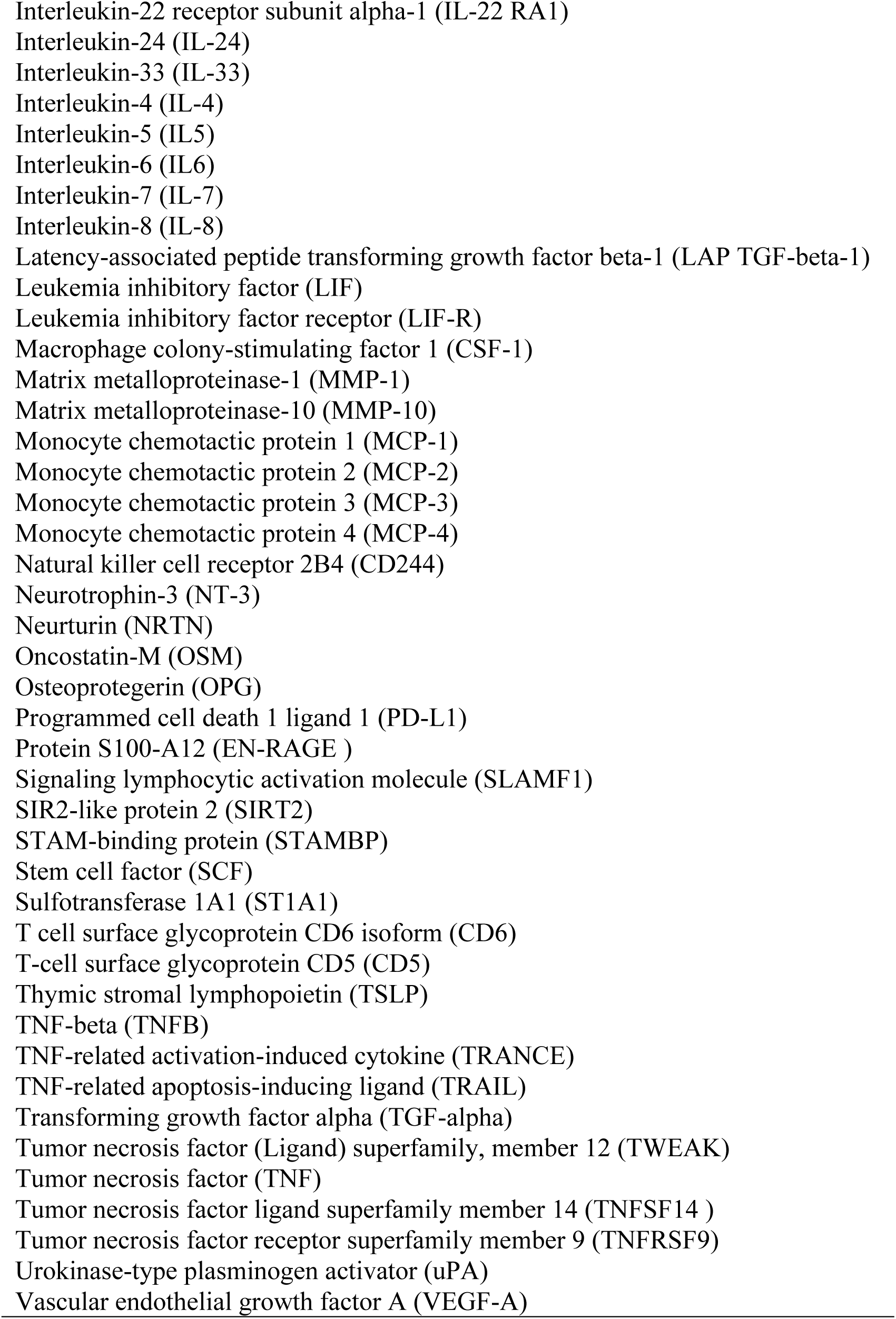
List of biomarkers in Olink’s inflammation panel.

**Table S4:** The percentage of plasma samples from CBD and patients groups together that have expressed the particular cytokine/marker.

|  | % Expression |
| --- | --- |
| **MMP-1** | 100 |
| **MCP-1** | 100 |
| **CD40** | 100 |
| **uPA** | 100 |
| **SCF** | 100 |
| **OPG** | 100 |
| **VEGF-A** | 100 |
| **CXCL5** | 100 |
| **CCL23** | 100 |
| **TWEAK** | 100 |
| **CCL19** | 100 |
| **Flt3L** | 100 |
| **CCL11** | 100 |
| **IL-18** | 100 |
| **MCP-2** | 100 |
| **CXCL11** | 100 |
| **CXCL1** | 100 |
| **TRAIL** | 100 |
| **CXCL10** | 100 |
| **DNER** | 100 |
| **FGF-19** | 100 |
| **CSF-1** | 100 |
| **CXCL6** | 100 |
| **HGF** | 100 |
| **4E-BP1** | 100 |
| **IL-18R1** | 100 |
| **CCL4** | 100 |
| **CXCL9** | 100 |
| **LAP TGF-beta-1** | 100 |
| **IL-10RB** | 100 |
| **IL-10RA** | 33 |
| **IL-8** | 100 |
| **CST5** | 100 |
| **CD244** | 100 |
| **TNFRSF9** | 100 |
| **MMP-10** | 100 |
| **CD5** | 100 |
| **AXIN1** | 100 |
| **CCL25** | 100 |
| **CX3CL1** | 100 |
| **IL-12B** | 100 |
| **FGF-21** | 100 |
| **STAMPB** | 100 |
| **TRANCE** | 100 |
| **CD6** | 100 |
| **CCL20** | 100 |
| **ADA** | 100 |
| **TNFSF14** | 100 |
| **TNFB** | 100 |
| **LIF-R** | 100 |
| **SIRT2** | 96 |
| **PD-L1** | 100 |
| **MIP-1 alpha** | 100 |
| **IL-6** | 43 |
| **IL-7** | 100 |
| **OSM** | 100 |
| **ST1A1** | 88 |
| **MCP-4** | 100 |
| **CASP-8** | 96 |
| **SLAMF1** | 86 |
| **TGF-alpha** | 100 |
| **FGF-23** | 100 |
| **CDCP1** | 100 |
| **Beta-NGF** | 84 |
| **CCL28** | 100 |
| **EN-RAGE** | 88 |
| **NT-3** | 70 |

**Table S5:**
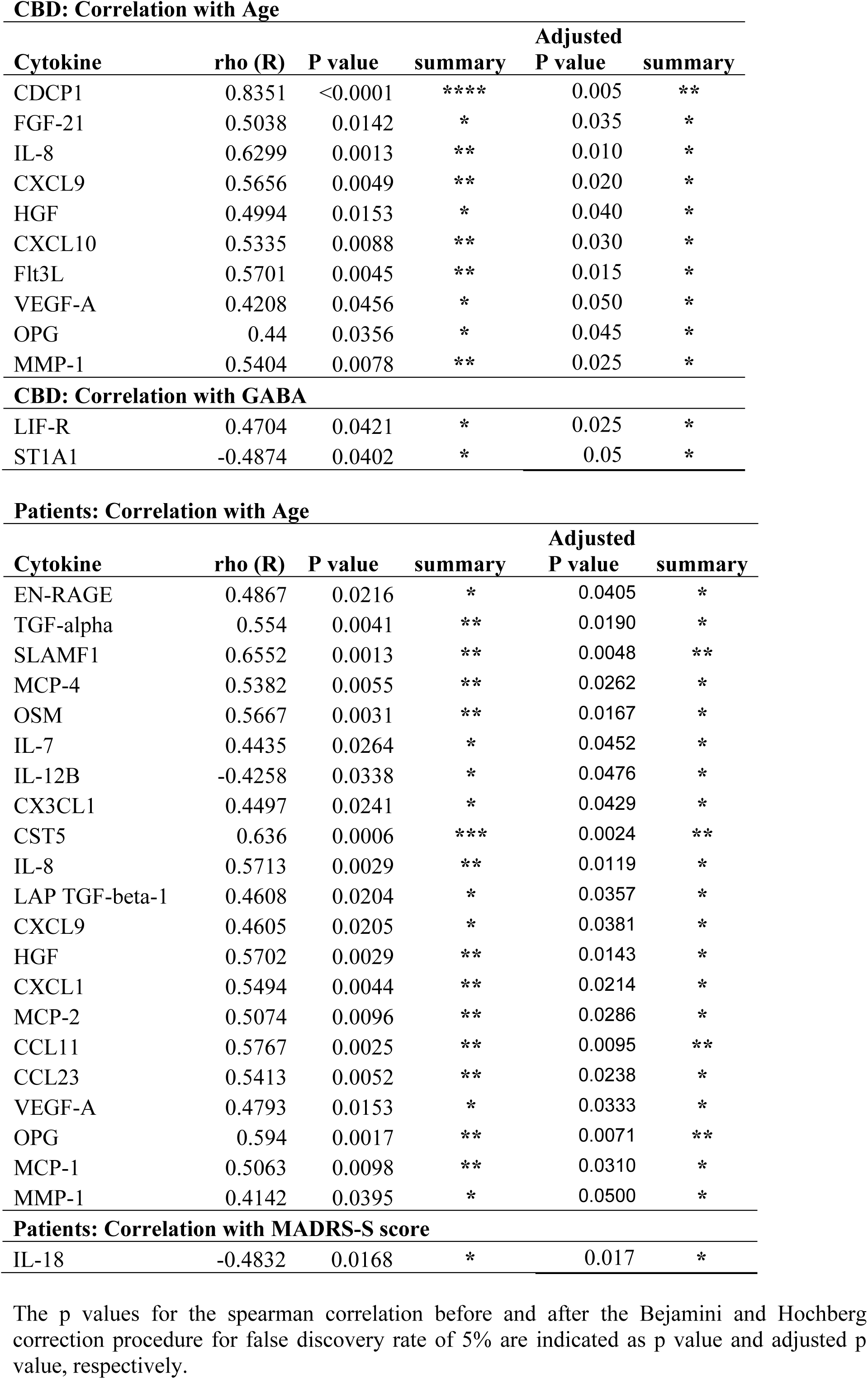
Correlation of expression level of cytokines in plasma with age or GABA concentration in CBD and patients.

**Table S6:** The percentage of supernatant samples of stimulated PBMC from psychiatric patients that have expressed the particular cytokine/marker.

|  | **% Expression** |
| --- | --- |
| **IL-8** | 100 |
| **TNF** | 100 |
| **CCL4** | 100 |
| **CXCL10** | 100 |
| **MMP-1** | 100 |
| **MCP-1** | 100 |
| **CXCL1** | 100 |
| **MCP-2** | 100 |
| **CXCL5** | 100 |
| **MIP-1 alpha** | 100 |
| **TNFRSF9** | 100 |
| **OSM** | 100 |
| **TNFB** | 100 |
| **CXCL9** | 100 |
| **LIF** | 100 |
| **IL-17A** | 100 |
| **IL-6** | 100 |
| **CCL20** | 100 |
| **IFN-gamma** | 100 |
| **4E-BP1** | 100 |
| **IL-1 alpha** | 100 |
| **CD40** | 100 |
| **MCP-3** | 100 |
| **uPA** | 100 |
| **CSF-1** | 100 |
| **TNFSF14** | 100 |
| **IL-12B** | 100 |
| **IL-13** | 100 |
| **CD5** | 100 |
| **Flt3L** | 100 |
| **IL-10** | 100 |
| **CASP-8** | 100 |
| **ADA** | 100 |
| **CXCL6** | 100 |
| **IL-18R1** | 100 |
| **CCL19** | 100 |
| **TRANCE** | 100 |
| **MMP-10** | 100 |
| **LAP TGF-beta-1** | 100 |
| **OPG** | 100 |
| **IL-2** | 86 |
| **STAMPB** | 100 |
| **PD-L1** | 100 |
| **VEGF-A** | 100 |
| **CD6** | 100 |
| **TWEAK** | 100 |
| **CXCL11** | 100 |
| **TGF-alpha** | 100 |
| **TRAIL** | 100 |
| **HGF** | 100 |
| **IL-4** | 100 |
| **SIRT2** | 100 |
| **IL-24** | 93 |
| **EN-RAGE** | 100 |
| **CCL23** | 100 |
| **IL-18** | 100 |
| **IL-5** | 83 |
| **CD244** | 100 |
| **DNER** | 100 |
| **CDCP1** | 100 |
| **AXIN1** | 100 |
| **SLAMF1** | 89 |
| **SCF** | 100 |
| **IL-10RA** | 85 |
| **ST1A1** | 100 |
| **IL-2RB** | 63 |
| **IL-15RA** | 70 |
| **CCL28** | 61 |

**Table S7:** GABA treatment alters cytokines released by PBMCs from patients.

| **GABA 100 nM** |  |
| --- | --- |
| **Markers** | **P values** |
| **AXIN1** | 0.016 |
| **TNFRSF9** | 0.015 |
| **VEGF-A** | 0.016 |
| **IL-1α** | 0.015 |

| **GABA 500 nM** |  |
| --- | --- |
| **Markers** | **P values** |
| **CD244** | 0.016 |
| **IL-13** | 0.016 |
| **HGF** | 0.015 |
| **VEGF-A** | 0.003 |

**Table S8:** Shapiro-Wilk normality test.

|  | P value | Passed<br>normality test? | Test performed |
| --- | --- | --- | --- |
| **GABA receptor subunits** |  |  |  |
| ρ2 | <0.0001 | No | Mann-Whitney |
| GABA-B1 | 0.0027 | No | Mann-Whitney |
| **Chloride transporters** |  |  |  |
| NKCC1 | 0.088 | Yes | Unpaired T-test |
| KCC1 | <0.0001 | No | Mann-Whitney |
| KCC3 | 0.0004 | No | Mann-Whitney |
| KCC4 | <0.0001 | No | Mann-Whitney |

## References

1. Olsen RW, Sieghart W (2008) International Union of Pharmacology. LXX. Subtypes of gamma-aminobutyric acid(A) receptors: classification on the basis of subunit composition, pharmacology, and function. Update. Pharmacol Rev 60: 243–60

2. Levite M (2008) Neurotransmitters activate T-cells and elicit crucial functions via neurotransmitter receptors. Curr Opin Pharmacol 8: 460–71

3. Tian J, Lu Y, Zhang H, Chau CH, Dang HN, Kaufman DL (2004) Gamma-aminobutyric acid inhibits T cell autoimmunity and the development of inflammatory responses in a mouse type 1 diabetes model. J Immunol 173: 5298–304

4. Bjurstom H, Wang J, Ericsson I, Bengtsson M, Liu Y, Kumar-Mendu S, Issazadeh-Navikas S, Birnir B (2008) GABA, a natural immunomodulator of T lymphocytes. J Neuroimmunol 205: 44–50

5. Bhat R, Axtell R, Mitra A, Miranda M, Lock C, Tsien RW, Steinman L (2010) Inhibitory role for GABA in autoimmune inflammation. Proc Natl Acad Sci U S A 107: 2580–5

6. Jin Z, Mendu SK, Birnir B (2013) GABA is an effective immunomodulatory molecule. Amino Acids 45: 87–94

7. Bhandage AK, Jin Z, Korol SV, Shen Q, Pei Y, Deng Q, Espes D, Carlsson PO, Kamali-Moghaddam M, Birnir B (2018) GABA Regulates Release of Inflammatory Cytokines From Peripheral Blood Mononuclear Cells and CD4(+) T Cells and Is Immunosuppressive in Type 1 Diabetes. EBioMedicine 30: 283–294

8. Miller AH, Raison CL (2016) The role of inflammation in depression: from evolutionary imperative to modern treatment target. Nat Rev Immunol 16: 22–34

9. Dantzer R (2001) Cytokine-induced sickness behavior: mechanisms and implications. Ann N Y Acad Sci 933: 222–34

10. Udina M, Castellvi P, Moreno-Espana J, Navines R, Valdes M, Forns X, Langohr K, Sola R, Vieta E, Martin-Santos R (2012) Interferon-induced depression in chronic hepatitis C: a systematic review and meta-analysis. J Clin Psychiatry 73: 1128–38

11. Dowlati Y, Herrmann N, Swardfager W, Liu H, Sham L, Reim EK, Lanctot KL (2010) A meta-analysis of cytokines in major depression. Biol Psychiatry 67: 446–57

12. Miller AH, Maletic V, Raison CL (2009) Inflammation and its discontents: the role of cytokines in the pathophysiology of major depression. Biol Psychiatry 65: 732–41

13. Jones KA, Thomsen C (2013) The role of the innate immune system in psychiatric disorders. Mol Cell Neurosci 53: 52–62

14. Valkanova V, Ebmeier KP, Allan CL (2013) CRP, IL-6 and depression: a systematic review and meta-analysis of longitudinal studies. J Affect Disord 150: 736–44

15. Raison CL, Rutherford RE, Woolwine BJ, Shuo C, Schettler P, Drake DF, Haroon E, Miller AH (2013) A randomized controlled trial of the tumor necrosis factor antagonist infliximab for treatment-resistant depression: the role of baseline inflammatory biomarkers. JAMA Psychiatry 70: 31–41

16. Strawbridge R, Arnone D, Danese A, Papadopoulos A, Herane Vives A, Cleare AJ (2015) Inflammation and clinical response to treatment in depression: A meta-analysis. Eur Neuropsychopharmacol 25: 1532–43

17. Kohler CA, Freitas TH, Stubbs B, Maes M, Solmi M, Veronese N, de Andrade NQ, Morris G, Fernandes BS, Brunoni AR, et al. (2017) Peripheral Alterations in Cytokine and Chemokine Levels After Antidepressant Drug Treatment for Major Depressive Disorder: Systematic Review and Meta-Analysis. Mol Neurobiol

18. Kohler CA, Freitas TH, Maes M, de Andrade NQ, Liu CS, Fernandes BS, Stubbs B, Solmi M, Veronese N, Herrmann N, et al. (2017) Peripheral cytokine and chemokine alterations in depression: a meta-analysis of 82 studies. Acta Psychiatr Scand 135: 373–387

19. Benedetti F, Poletti S, Hoogenboezem TA, Locatelli C, de Wit H, Wijkhuijs AJM, Colombo C, Drexhage HA (2017) Higher Baseline Proinflammatory Cytokines Mark Poor Antidepressant Response in Bipolar Disorder. J Clin Psychiatry 78: e986–e993

20. Uher R, Tansey KE, Dew T, Maier W, Mors O, Hauser J, Dernovsek MZ, Henigsberg N, Souery D, Farmer A, et al. (2014) An inflammatory biomarker as a differential predictor of outcome of depression treatment with escitalopram and nortriptyline. Am J Psychiatry 171: 1278–86

21. Cattaneo A, Gennarelli M, Uher R, Breen G, Farmer A, Aitchison KJ, Craig IW, Anacker C, Zunsztain PA, McGuffin P, et al. (2013) Candidate genes expression profile associated with antidepressants response in the GENDEP study: differentiating between baseline ‘predictors’ and longitudinal ‘targets’. Neuropsychopharmacology 38: 377–85

22. Bu DF, Erlander MG, Hitz BC, Tillakaratne NJ, Kaufman DL, Wagner-McPherson CB, Evans GA, Tobin AJ (1992) Two human glutamate decarboxylases, 65-kDa GAD and 67-kDa GAD, are each encoded by a single gene. Proc Natl Acad Sci U S A 89: 2115–9

23. Schmidt MJ, Mirnics K (2015) Neurodevelopment, GABA system dysfunction, and schizophrenia. Neuropsychopharmacology 40: 190–206

24. Korol SV JZ, Jin Y, Bhandage AK, Tengholm A, Gandasi N, Barg S, Espes D, Carlsson PO, Laver D, Birnir B (2018) Functional characterization of native, high-affinity GABAA receptors in human pancreatic beta cells eBioMedicine Accepted for publication: 273–282

25. Birnir B, Korpi ER (2007) The impact of sub-cellular location and intracellular neuronal proteins on properties of GABA(A) receptors. Curr Pharm Des 13: 3169–77

26. Petty F, Sherman AD (1984) Plasma GABA levels in psychiatric illness. J Affect Disord 6: 131–8

27. Plog BA, Nedergaard M (2018) The Glymphatic System in Central Nervous System Health and Disease: Past, Present, and Future. Annu Rev Pathol 13: 379–394

28. Bhandage AK, Hellgren C, Jin Z, Olafsson EB, Sundstrom-Poromaa I, Birnir B (2015) Expression of GABA receptors subunits in peripheral blood mononuclear cells is gender dependent, altered in pregnancy and modified by mental health. Acta Physiol (Oxf) 213: 575–85

29. Conze T, Lammers R, Kuci S, Scherl-Mostageer M, Schweifer N, Kanz L, Buhring HJ (2003) CDCP1 is a novel marker for hematopoietic stem cells. Ann N Y Acad Sci 996: 222–6

30. Spassov DS, Wong CH, Sergina N, Ahuja D, Fried M, Sheppard D, Moasser MM (2011) Phosphorylation of Trask by Src kinases inhibits integrin clustering and functions in exclusion with focal adhesion signaling. Mol Cell Biol 31: 766–82

31. Enyindah-Asonye G, Li Y, Ruth JH, Spassov DS, Hebron KE, Zijlstra A, Moasser MM, Wang B, Singer NG, Cui H, et al. (2017) CD318 is a ligand for CD6. Proc Natl Acad Sci U S A 114: E6912–E6921

32. Hill LJ, Di Pietro V, Hazeldine J, Davies D, Toman E, Logan A, Belli A (2017) Cystatin D (CST5): An ultra-early inflammatory biomarker of traumatic brain injury. Sci Rep 7: 5002

33. Ferrer-Mayorga G, Alvarez-Diaz S, Valle N, De Las Rivas J, Mendes M, Barderas R, Canals F, Tapia O, Casal JI, Lafarga M, et al. (2015) Cystatin D locates in the nucleus at sites of active transcription and modulates gene and protein expression. J Biol Chem 290: 26533–48

34. Marshall FH, Jones KA, Kaupmann K, Bettler B (1999) GABAB receptors - the first 7TM heterodimers. Trends Pharmacol Sci 20: 396–9

35. Olsen RW, Sieghart W (2009) GABA A receptors: subtypes provide diversity of function and pharmacology. Neuropharmacology 56: 141–8

36. Gassmann M, Shaban H, Vigot R, Sansig G, Haller C, Barbieri S, Humeau Y, Schuler V, Muller M, Kinzel B, et al. (2004) Redistribution of GABAB(1) protein and atypical GABAB responses in GABAB(2)-deficient mice. J Neurosci 24: 6086–97

37. Pallavi P, Sagar R, Mehta M, Sharma S, Subramanium A, Shamshi F, Sengupta U, Qadri R, Pandey RM, Mukhopadhyay AK (2013) Serum neurotrophic factors in adolescent depression: gender difference and correlation with clinical severity. J Affect Disord 150: 415–23

38. Oglodek EA, Just MJ, Szromek AR, Araszkiewicz A (2016) Melatonin and neurotrophins NT-3, BDNF, NGF in patients with varying levels of depression severity. Pharmacol Rep 68: 945–51

39. Quiroz JA, Machado-Vieira R, Zarate CA, Jr., Manji HK (2010) Novel insights into lithium’s mechanism of action: neurotrophic and neuroprotective effects. Neuropsychobiology 62: 50–60

40. Engel D, Zomkowski AD, Lieberknecht V, Rodrigues AL, Gabilan NH (2013) Chronic administration of duloxetine and mirtazapine downregulates proapoptotic proteins and upregulates neurotrophin gene expression in the hippocampus and cerebral cortex of mice. J Psychiatr Res 47: 802–8

41. Angelucci F, Aloe L, Jimenez-Vasquez P, Mathe AA (2003) Lithium treatment alters brain concentrations of nerve growth factor, brain-derived neurotrophic factor and glial cell line-derived neurotrophic factor in a rat model of depression. Int J Neuropsychopharmacol 6: 225–31

42. Hellweg R, Lang UE, Nagel M, Baumgartner A (2002) Subchronic treatment with lithium increases nerve growth factor content in distinct brain regions of adult rats. Mol Psychiatry 7: 604–8

43. Walz JC, Frey BN, Andreazza AC, Cereser KM, Cacilhas AA, Valvassori SS, Quevedo J, Kapczinski F (2008) Effects of lithium and valproate on serum and hippocampal neurotrophin-3 levels in an animal model of mania. J Psychiatr Res 42: 416–21

44. Hannestad J, DellaGioia N, Bloch M (2011) The effect of antidepressant medication treatment on serum levels of inflammatory cytokines: a meta-analysis. Neuropsychopharmacology 36: 2452–9

45. Jansen R, Penninx BW, Madar V, Xia K, Milaneschi Y, Hottenga JJ, Hammerschlag AR, Beekman A, van der Wee N, Smit JH, et al. (2016) Gene expression in major depressive disorder. Mol Psychiatry 21: 339–47

46. Svenningsson P, Berg L, Matthews D, Ionescu DF, Richards EM, Niciu MJ, Malinger A, Toups M, Manji H, Trivedi MH, et al. (2014) Preliminary evidence that early reduction in p11 levels in natural killer cells and monocytes predicts the likelihood of antidepressant response to chronic citalopram. Mol Psychiatry 19: 962–4

47. Ryan KM, McLoughlin DM (2018) Vascular endothelial growth factor plasma levels in depression and following electroconvulsive therapy. Eur Arch Psychiatry Clin Neurosci 268: 839–848

48. Maes M (2001) The immunoregulatory effects of antidepressants. Hum Psychopharmacol 16: 95–103

49. Gadad BS, Jha MK, Grannemann BD, Mayes TL, Trivedi MH (2017) Proteomics profiling reveals inflammatory biomarkers of antidepressant treatment response: Findings from the CO-MED trial. J Psychiatr Res 94: 1–6

50. Ricken R, Busche M, Schlattmann P, Himmerich H, Bopp S, Bschor T, Richter C, Stamm TJ, Heinz A, Hellweg R, et al. (2018) Cytokine serum levels remain unchanged during lithium augmentation of antidepressants in major depression. J Psychiatr Res 96: 203–208

51. Leighton SP, Nerurkar L, Krishnadas R, Johnman C, Graham GJ, Cavanagh J (2018) Chemokines in depression in health and in inflammatory illness: a systematic review and meta-analysis. Mol Psychiatry 23: 48–58

52. Ho PS, Yeh YW, Huang SY, Liang CS (2015) A shift toward T helper 2 responses and an increase in modulators of innate immunity in depressed patients treated with escitalopram. Psychoneuroendocrinology 53: 246–55

53. Fan N, Luo Y, Ou Y, He H (2017) Altered serum levels of TNF-alpha, IL-6, and IL-18 in depressive disorder patients. Hum Psychopharmacol 32

54. Brann E, Fransson E, White RA, Papadopoulos FC, Edvinsson A, Kamali-Moghaddam M, Cunningham JL, Sundstrom-Poromaa I, Skalkidou A (2018) Inflammatory markers in women with postpartum depressive symptoms. J Neurosci Res

55. Herder C, Hermanns N (2019) Subclinical inflammation and depressive symptoms in patients with type 1 and type 2 diabetes. Semin Immunopathol

56. Pietrzak D, Pietrzak A, Grywalska E, Kicinski P, Rolinski J, Donica H, Franciszkiewicz-Pietrzak K, Borzecki A, Socha M, Niedzialek J, et al. (2018) Serum concentrations of interleukin 18 and 25-hydroxyvitamin D3 correlate with depression severity in men with psoriasis. PLoS One 13: e0201589

57. Antonelli A, Rotondi M, Fallahi P, Ferrari SM, Paolicchi A, Romagnani P, Serio M, Ferrannini E (2006) Increase of CXC chemokine CXCL10 and CC chemokine CCL2 serum levels in normal ageing. Cytokine 34: 32–8

58. Enroth S, Enroth SB, Johansson A, Gyllensten U (2015) Protein profiling reveals consequences of lifestyle choices on predicted biological aging. Sci Rep 5: 17282

59. Larsson A, Carlsson L, Gordh T, Lind AL, Thulin M, Kamali-Moghaddam M (2015) The effects of age and gender on plasma levels of 63 cytokines. J Immunol Methods 425: 58–61

60. Shurin GV, Yurkovetsky ZR, Chatta GS, Tourkova IL, Shurin MR, Lokshin AE (2007) Dynamic alteration of soluble serum biomarkers in healthy aging. Cytokine 39: 123–9

61. Petty F (1995) GABA and mood disorders: a brief review and hypothesis. J Affect Disord 34: 275–81

62. Petty F, Kramer GL, Dunnam D, Rush AJ (1990) Plasma GABA in mood disorders. Psychopharmacol Bull 26: 157–61

63. Petty F, Kramer GL, Fulton M, Moeller FG, Rush AJ (1993) Low plasma GABA is a trait-like marker for bipolar illness. Neuropsychopharmacology 9: 125–32

64. Petty F, Schlesser MA (1981) Plasma GABA in affective illness. A preliminary investigation. J Affect Disord 3: 339–43

65. Berrettini WH, Nurnberger JI, Jr., Hare TA, Simmons-Alling S, Gershon ES, Post RM (1983) Reduced plasma and CSF gamma-aminobutyric acid in affective illness: effect of lithium carbonate. Biol Psychiatry 18: 185–94

66. Lu YR, Fu XY, Shi LG, Jiang Y, Wu JL, Weng XJ, Wang ZP, Wu XY, Lin Z, Liu WB, et al. (2014) Decreased plasma neuroactive amino acids and increased nitric oxide levels in melancholic major depressive disorder. BMC Psychiatry 14: 123

67. Loscher W (1982) Anticonvulsant and biochemical effects of inhibitors of GABA aminotransferase and valproic acid during subchronic treatment in mice. Biochem Pharmacol 31: 837–42

68. Barragan A, Weidner JM, Jin Z, Korpi ER, Birnir B (2015) GABAergic signalling in the immune system. Acta Physiol (Oxf) 213: 819–27

69. Fuks JM, Arrighi RB, Weidner JM, Kumar Mendu S, Jin Z, Wallin RP, Rethi B, Birnir B, Barragan A (2012) GABAergic signaling is linked to a hypermigratory phenotype in dendritic cells infected by Toxoplasma gondii. PLoS Pathog 8: e1003051

70. Sheehan DV, Lecrubier Y, Sheehan KH, Amorim P, Janavs J, Weiller E, Hergueta T, Baker R, Dunbar GC (1998) The Mini-International Neuropsychiatric Interview (M.I.N.I.): the development and validation of a structured diagnostic psychiatric interview for DSM-IV and ICD-10. J Clin Psychiatry 59 Suppl 20: 22–33;quiz 34-57

71. Svanborg P, Asberg M (1994) A new self-rating scale for depression and anxiety states based on the Comprehensive Psychopathological Rating Scale. Acta Psychiatr Scand 89: 21–8

72. Cunningham JL, Wernroth L, von Knorring L, Berglund L, Ekselius L (2011) Agreement between physicians’ and patients’ ratings on the Montgomery-Asberg Depression Rating Scale. J Affect Disord 135: 148–53

73. Edvinsson A, Brann E, Hellgren C, Freyhult E, White R, Kamali-Moghaddam M, Olivier J, Bergquist J, Bostrom AE, Schioth HB, et al. (2017) Lower inflammatory markers in women with antenatal depression brings the M1/M2 balance into focus from a new direction. Psychoneuroendocrinology 80: 15–25

74. Benjamini Y, Hochberg Y (1995) Controlling the False Discovery Rate - a Practical and Powerful Approach to Multiple Testing. J R Stat Soc B 57: 289–300

